# High-Quality PacBio Genome Assembly of *Populus alba* L. “Villafranca”

**DOI:** 10.1101/2025.06.30.661909

**Authors:** Iqra Sarfraz, Andrea Zuccolo, Mirko Celii, Alessandra Francini, Rod A. Wing, Luca Sebastiani

**Affiliations:** Institute of Crop Science, Scuola Superiore Sant’Anna, Piazza Martiri della Libertà 33, 56127, Pisa; King Abdullah University of Science & Technology (KAUST), Saudi Arabia; Arizona Genomics Institute, School of Plant Sciences, University of Arizona, Tucson, AZ, USA

## Abstract

This study presents the high-quality genome assemblies for *Populus alba* L. “Villafranca” using PacBio HiFi sequencing. The assembly span 498.95 Mb, an N50 of 18.18 Mb and largest contig of 52.03 Mb. BUSCO analysis revealed genome completeness (embryophyta_odb10) with 98.8% of the 1,614 BUSCO groups searched. The Transposable element and repetitive content accounted for ∼31.37%. The comparison of *P. alba* and *P. trichocarpa* genomes identified 9,741 structural variants (SVs) This comprehensive analysis provides valuable resources for studying poplar genome evolution, domestication, and genetic improvement, underscoring the utility of long-read sequencing for resolving complex genomic features.

## Background & Summary

Poplar plants are part of the genus *Populus*, which belongs to the family *Salicaceae*. They possess many important qualities, such as their flexible adaptability, large biomass production, smooth propagation, hybridization tendency, relatively easy transformability, and the availability of protocols for *in vitro* cultivation and micro-propagation (1,2). Poplars are an important source of essential forest commodities, including wood, fuelwood, and fiber and play a significant role in the reintegration and renewal of degraded landscapes (3). Most notably, their vast carbon storage capacity is crucial for mitigating climate change (4).

The white poplar (*Populus alba*), also known as silver poplar, is a deciduous tree which is widely distributed across Europe, and in central Asia. *P. alba* is resistant to pests, fungal and bacterial pathogens (5). It is also known to tolerate various abiotic stresses such as drought, salinity, heavy metals and low temperatures and it plays an important role in phytoremediation (6,7). Specifically, the “Villafranca” clone has been extensively studied in recent years, because of its valuable traits, such as tolerance to heavy metals like zinc (Zn), and cadmium (Cd) (6,8). Also, “Villafranca” clone has been successfully transformed using *Agrobacterium tumefaciens* mediated gene transfer (9). These characteristics make “Villafranca” an ideal candidate for genome level studies, which has been made possible due to advancements in sequencing technologies.

The Sanger based successful sequencing and assembly of human genome in 2001, marked a turning point, paving the way for the study of larger genomes (10). Before the introduction of next generation sequencing (NGS), genomics primarily centred on the characterization and identification of single genes due to limitation of classical sequencing techniques.

One of the most notable successes of these early genome sequencing effort was the sequencing of *Populus trichocarpa*, which is native to western North America (11). This genome was assembled following a whole genome shotgun sequencing, revealing an estimated genome size of approximately 485 megabases (Mb) and nearly 45,000 protein coding genes (12). This reference assembly provided a comprehensive framework for understanding genomic organization, identifying genetic variations, and facilitating functional genomic studies. However, early genome assemblies contained unresolved gaps, particularly in structurally complex and repetitive areas of genome. In the recent years the widespread adoption of Third generation sequencing technologies, such as PacBio, Oxford Nanopore have overcome these limitations being capable to generate long DNA reads (13). The use of these technologies enabled the generation of an improved version of the *P. trichocarpa* assembly (GCF_000002775.5), allowing better identification of structural variants (SVs), transposable elements (TEs) and repeats (14).

A draft genome assembly of *P. alba* based on PacBio RSII and Illumina technology was released in 2019 by Chinese academy of forestry. This draft assembly, covered genome size of 416 Mb and included 30,624 protein coding genes, Liu et al. (15). The most recent assembly of *P. alba* submitted by the Chinese academy of forestry was based on Illumina sequencing and Hi-C data, covers 416 .4 Mb, with scaffold N50 of 22.7 Mb, A contig N50 of 1.2 Mb, GC% of 33.5%, and included a total 34,010 protein coding genes (16). These assemblies provide a valuable genomic resource for P. alba; however, they are based on the PAL-ZL1 isolate, a genotype from China that is geographically distinct from European populations. In contrast, our study focuses on generating a high-quality, long read genome assembly for the European isolaate *P. alba* L. “Villafranca” By using PacBio HiFi sequencing, this study generated a high-quality assembly of 498.95 Mb, an N50 of 18.18 Mb and a largest contig of 52.02 Mb. The genome comprises 44,852 predicted genes (71.9% functionally annotated), and 9,741 SVs relative to *P*.*trichocarpa* .This assembly allows better annotation of genes and improves the resolution in repetitive regions and facilitates more accurate detection of SVs, a key aspect of understanding stress adaptation mechanism in. This comprehensive analysis contributes to the expanding genomic resources for populus and allows further investigation into genome evolution, domestication, and genetic improvement related to climate resilience and stress response.

## Methods

### Plant material and growth conditions

*P. alba* L. *“*Villafranca*”* clone plantlets were grown *in vitro* in magenta vessels on a woody plant medium (WPM) at pH 5.7 (17) and root development medium (RD11) in a controlled environment room (22 °C temperature and 16:8h light cycle period). Plantlets were propagated through *in vitro* cultures of apexes and nodal explants.

### Genomic DNA extraction and Sequencing

Young leaves of *P. alba* L. “Villafranca”, preserved in liquid nitrogen and stored at -80 °C were used for the extraction of High-quality DNA using a modified CTAB protocol (18). The quality of the extracted DNA was assessed through pulsed-field gel electrophoresis (CHEF) on 1% agarose gels to evaluate fragment size and restriction enzyme digestibility. Quantification was performed using Qubit fluorometry (Thermo Fisher Scientific, Waltham, MA). Sequencing was performed using PacBio Revio System at the Arizona Genomics Institute (Tucson, AZ), on Revio SMRT cells (19). A total of 35.02 Gbp of HiFi reads for *P. alba* L. “Villafranca” were generated, with N50 value of 17.07 kbp.

### Genome size and heterozygosity estimation

To estimate genome size and heterozygosity, reads obtained from PacBio sequencing were subjected to 21-mer frequency distribution using the efficient and fast k-mer counting software Jellyfish v2.3.0 (20). The results of the analysis were then processed using Genomescope2 (21). The K-mer distribution displayed two distinct peaks, one at approximately 37.8x coverage corresponds to heterozygous region while, the second peak at around 76x coverage represents the homozygous region. The heterozygosity rate was estimated to be 1.17%, indicating the genome is predominantly homozygous. The duplication rate was found to be 0.25% suggesting a low level of duplicated content. Approximately, 66.9% of genome was composed of unique sequences with remaining 33% being repetitive (Fig 1).

**Figure 1.**
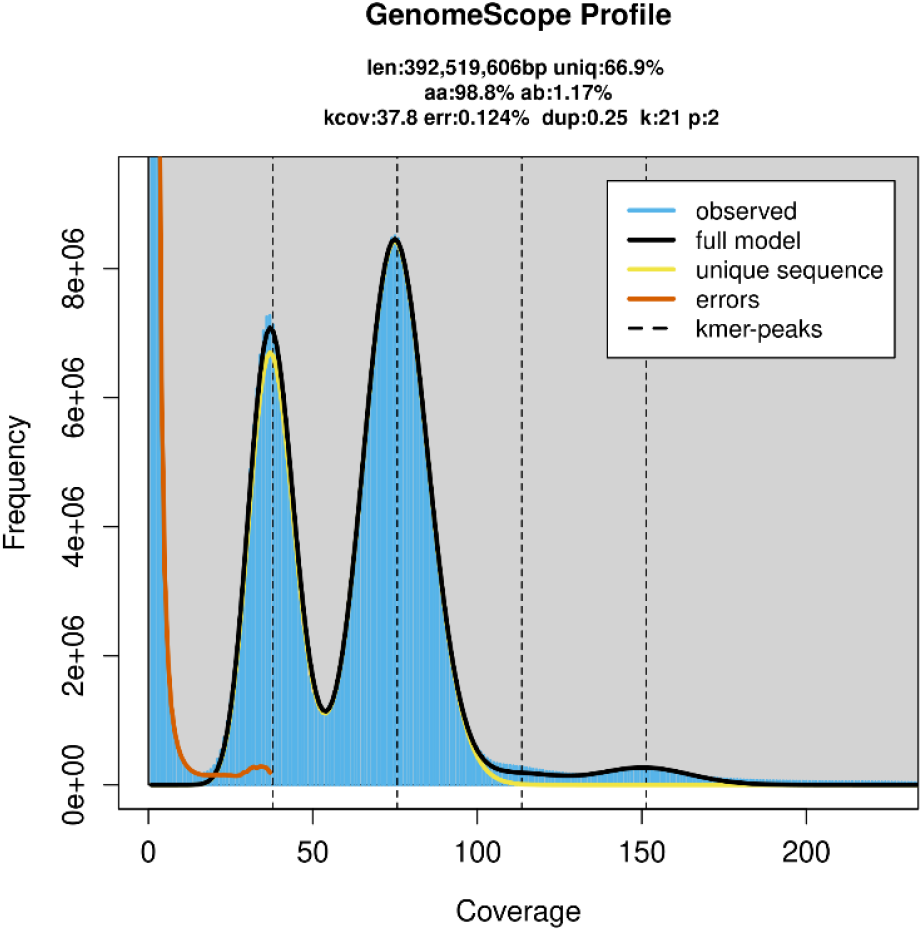
Genome size estimation generated using GenomeScope. Using a K-mer analysis (K=21, from jellyfish).

### Genome assembly and scaffolding

The PacBio HiFi reads were assembled using hifiasm/v0.19.8 (22) with default parameters. The redundant haplotigs were removed with the help of purge haplotigs (23). the quality assessment tool for genome assemblies QUAST/v5.2.0 (24) was used to evaluate primary assembly. The *P. alba* L. “Villafranca” assembly comprised of 925 contigs with a total length of 498.95 Mb, an N50 of 18.18 Mb and largest contig of 52.03 Mb. The GC content of assembly was 36.44%.

**Table 1.** Main statistics of *P. alba* L. “Villafranca” draft genome assembly.

| Parameter | *P. alba* L.<br>“Villafranca” |
| --- | --- |
| Total number of contigs | 925 |
| Total number of contigs ($\geq 50,000$ bp) | 361 |
| Total length ($\geq 50,000$ bp) | 478,315,314 |
| Largest Contigs | 52,030,799 |
| GC % | 36.44 |
| N50 | 18,489,627 |
| N90 | 242,455 |
| L50 | 10 |
| L90 | 49 |

To assess the completeness of genome, Benchmarking Universal Single-Copy Orthologs (BUSCO)/v5.7.1 (25) was run in genome mode using the embryophyte genes database (embryophyta_odb10). The BUSCO results showed that 98.8% of the 1,614 BUSCO groups were present and complete (C), with 83.52% of genes being single-copy and 15.2% duplicated. A small fraction of BUSCOs were fragmented (F:0.4%) or missing (M:0.9%) (Fig. 2). These results are quite similar to *P. alba* PAL-ZL1 isolate from Liu et al. (15) (98.3%) .

**Figure 2.**
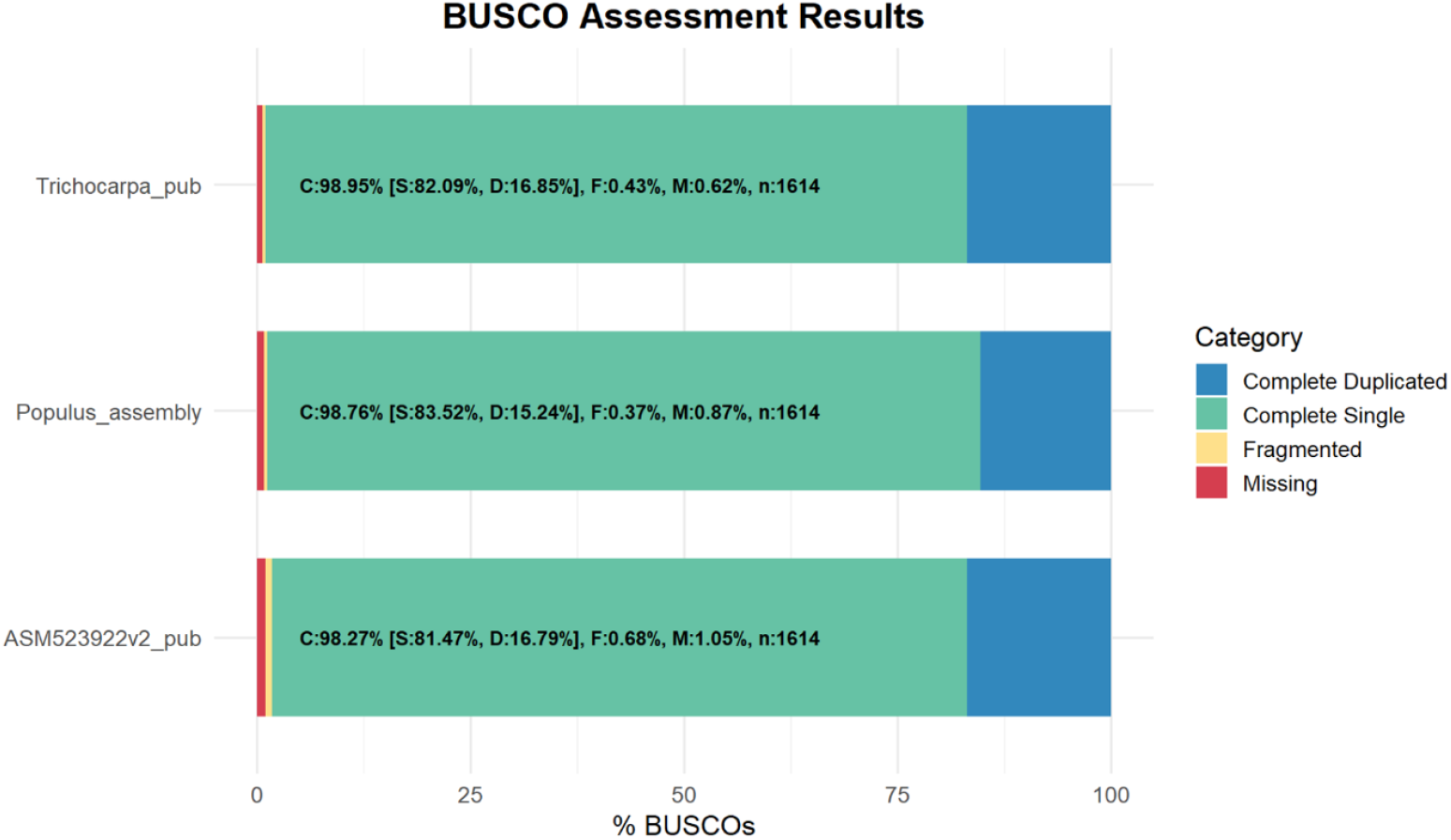
Comparison of genome completeness among different *Poplar* subsp. using BUSCO assessment. The figure demonstrates the percentage of complete (C), single (S), duplicated (D), fragmented (F) and missing (M) BUSCO orthologs identified in different genome assemblies. Total BUSCO genes n=1,614.

*P. tremula* and *P. tremuloides* assemblies recovered 95% BUSCO genes with 1.7% and 1.6 % of missing genes respectively (26). Additionally, *P. euphratica* with 95.6% complete BUSCO, 13.47% duplicated and *P. trichocarpa* with 97.6% complete BUSCO for the genome and 98.7% for the gene annotation, 17.71% duplicated reported by Zhang et al. (27), further highlights the differences. Altogether, the BUSCO statistics testify the high completeness and contiguity achieved in our genome assembly.

### RagTag scaffolding

RagTag scaffold/v2.1.0 (28) was utilized for homology-based scaffolding. For the whole-genome alignment, the Minimap2 (29) inbuilt Ragtag aligner was used. The genome assembly of *P. alba* clone “Villafranca” was processed using the RagTag scaffold pipeline utilizing as a reference the *P. trichocarpa* genome assembly GCF-000002775.5. Ragtag aligns contigs to the reference genome, facilitating their placement along chromosome, as well as identifying any misplaced or unplaced contigs (Fig. 3 a) (28). Gap filling strategy was employed to reduce number of gaps. PacBio HiFi raw reads were first assembled using Flye assembler/v2.9.1 (30) in an assembly with estimated genome size of 500 Mbp. The Flye genome assembly was then combined with the Hifiasm one to further refine the genome using RagTag patch tool. The fasta files of both flye and hifiasm generated assemblies were provided as an input, fill-only parameter was employed to specifically target the gap filling without changing the rest of assembly.

**Figure 3.**
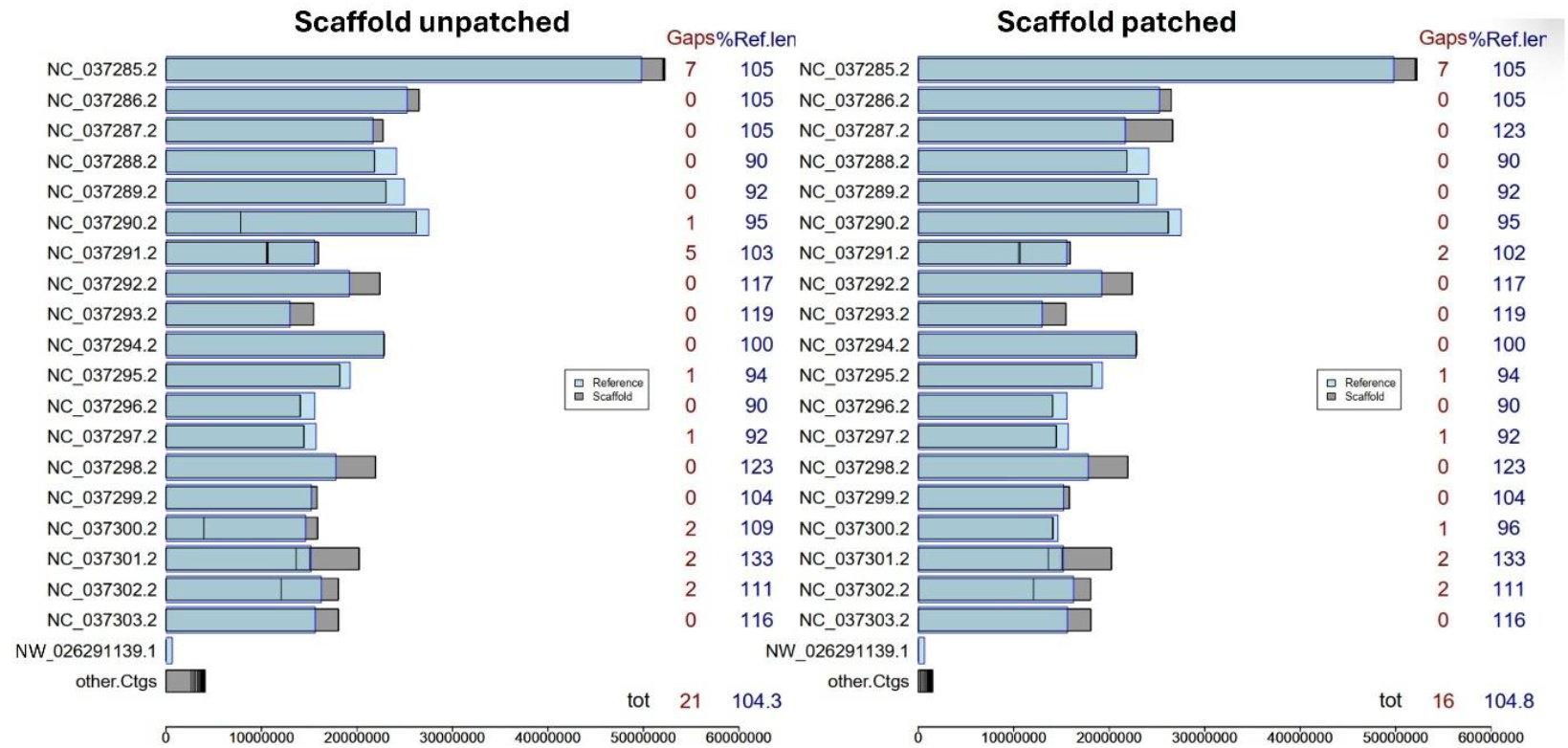
Comparison of *P. alba* L. “Villafranca” scaffolding before and after gap patching using *P. trichocarpa* as the reference genome. (a) Shows the scaffold length and percentage of reference genome coverage before the patching process (b) Shows the improvement in scaffold length and reduction in gaps after patching. Total number of gaps and the percentage of reference genome covered is given at the bottom of plot.

Scaffold alignment revealed notable structural discrepancies with P.trichocarpa reference, particularly on chromosome NC_037300.2 and NC-03728.2, which demonstrated large gaps and length differences, providing insight into potential rearrangement or assembly specific features. The gap filling strategy reduced the number of gaps reduced from 21 to 16 (Fig. 3b and 4).

**Figure 4.**
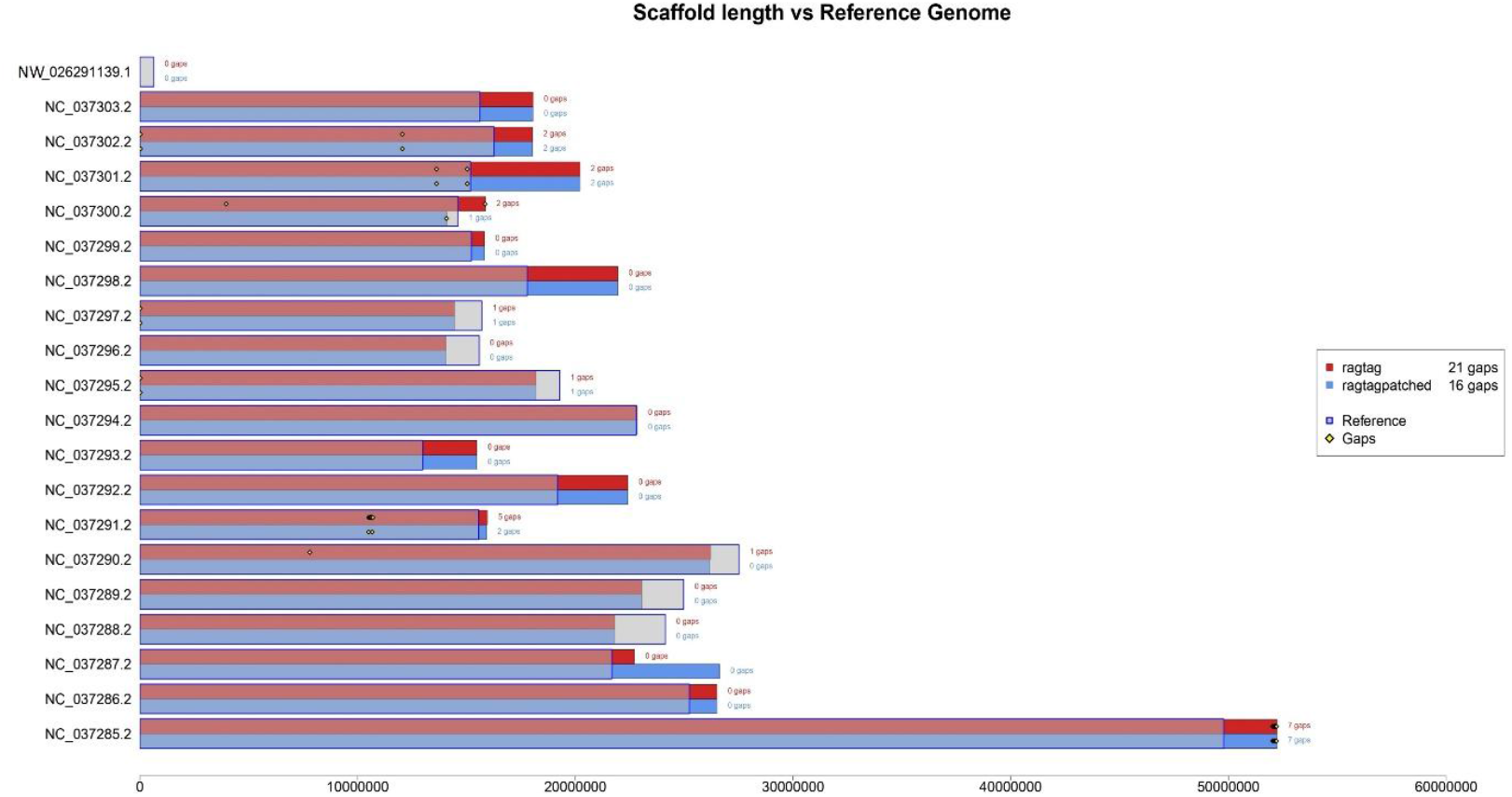
Scaffold length vs reference genome comparison for *P. alba* L. “Villafranca” scaffolding using *P. trichocarpa* as the reference genome. The plot compares scaffold lengths before and after gap patching. Reference genomes are represented by outlined blue bars. Gaps are marked with the diamond shapes on the bars.

### Transposable Element and Repeat annotation

To identify transposable elements (TEs) in the assembly the Extensive *de novo* TE annotator (EDTA)/v2.1.0 was used to produce a TE library(31). All the predicted Helitrons were removed from the TE library as their identification is not so imprecise and susceptible to generate false positives (31). The TE library was then used running RepeatMasker/v4.1.4 (32) under default settings onto our genome assembly. The resulting output file with masked TE sequences (soft masking) was used for the baseline annotation of genes.

The TE content of the draft assembly was estimated to be 31.37%, with retroelements accounting for 25.36%. Among these, long terminal repeats (LTR) were the most abundant, with Gypsy/DIRS1 elements alone covering 15.42% of the genome. DNA transposons represented 5.83% of the genome. A small fraction (0.18%) of unclassified were also observed (**Table** ).

**Table 2.** Transposable elements identified in *P. alba* L. clone “Villafranca”.

| Class | Number of elements | Length occupied | % of sequence |
| --- | --- | --- | --- |
| Retroelements | 177,936 | 126,513,305 bp | 25.36% |
| LTR elements: | 177,936 | 126,513,305 bp | 25.36% |
| Gypsy/DIRS1 | 57,399 | 76,927,293 bp | 15.42% |
| DNA transposons | 77,694 | 29,090,774 bp | 5.83% |
| Unclassified: | 6,195 | 899,557 bp | 0.18% |
| **Total interspersed repeats:** |  | 156,503,636 bp | 31.37% |

The total TE reported in *P. alba* L. “Villafranca” (31.37%) is lower than the repetitive content in genome of *P. alba* (45.16%) and *P. trichocarpa (*48.07%), as reported by Liu et al. (15). However, this discrepancy can be partially explained by methodological differences. Our analysis particularly focused on TEs and excluded Helitrons and microsatellites from the TEs library before repeat masking,. Liu et al. (15) included Helitrons and microsatellites in their analysis, which accounted for 4.17% and 4.08% respectively. After excluding these elements, the adjusted TE content in Liu et al. become 36.91 %. Additionally, our analysis did not include LINEs, which contributed another 1.22% in Liu’s dataset.

Therefore, the remaining 4% difference may reflect genuine variation in TE composition between Villafranca clone and *P*.*alba* genotype analyzed by Liu et al (15).

### Gene prediction and functional annotation

The gene prediction was carried out using Omicsbox (33), the eukaryotic gene finding module, which uses the AUGUSTUS algorithm (34). To enhance the accuracy of gene prediction model, a combination of ab-initio, homology-based and RNA-Seq supported approaches were utilized. *P. alba* RNA-Seq data with the accession numbers SRR15606815 and SRR15606816 were downloaded from NCBI Sequence Read Archive (SRA). Additionally, protein sequences from *P. trichocarpa* were used as a homologous reference in order to further support gene prediction process. The soft masked genome assembly was used as the primary input, with *Arabidopsis thaliana* gene model selected as the closest species.

The functional annotation was performed by comparing the predicted genes to the nr division of GeneBank using Diamond blast /v2.1.8 (35) and analyzing them with InterProScan/v5.61-93.0 (36). The parameters used for Diamond blast were Blast Mode = blastp, Sensitivity Mode = Standard, Database = NR, Blast e-value = 1.0 e^-3^. For InterProScan we used the Cloud InterProScan with up-to-date EMBL-EBI-InterPro data including CCD, HHM-Pfams and HMMPIR models. Afterwards, Gene Ontology (GO) mapping and annotation was carried out using the integrated pipeline available in omicsbox (33). Also, EggNOG-based functional annotation (37) was carried out within OmicsBox to assign orthologous groups. Merging EggNOG annotations with the initial GO and InterPro results, shows the increase in number of GO terms and enzyme codes , reflecting improved annotation quality shown in Fig. 5.

**Figure 5.**
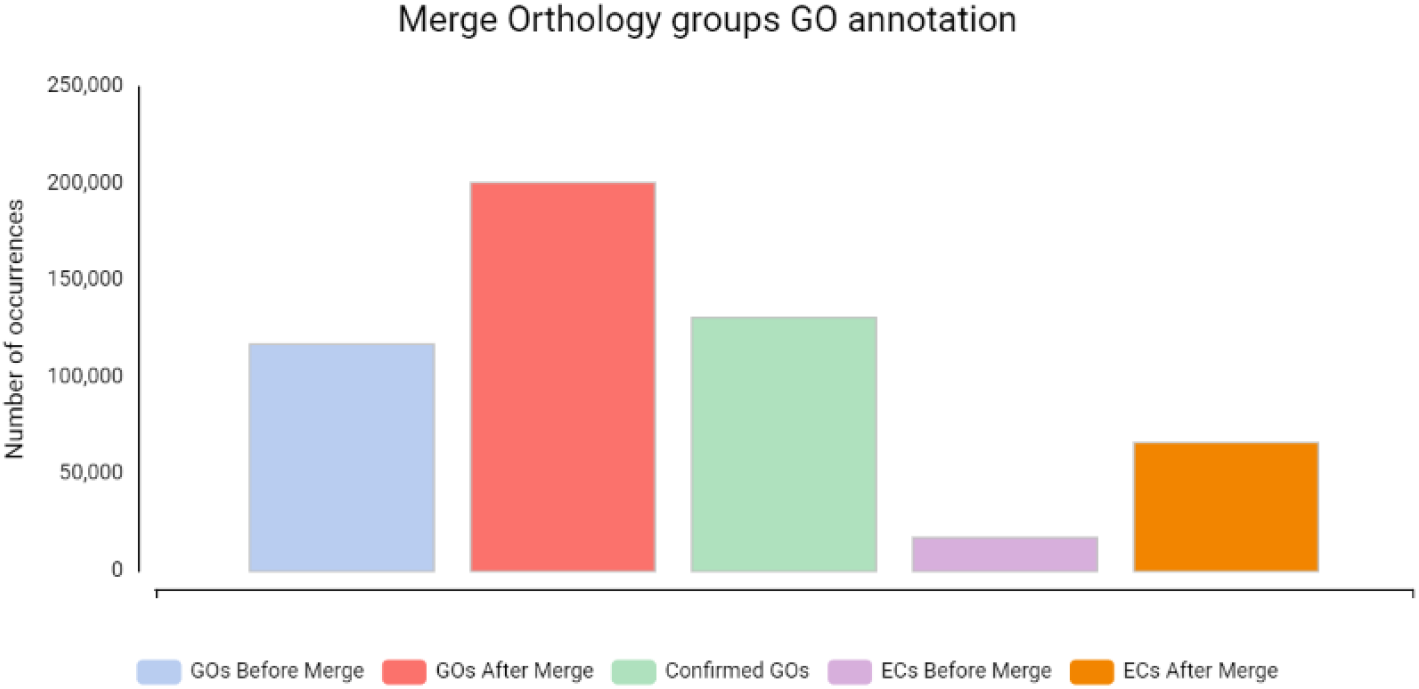
Improved GO and EC after combining EggNOG results with GO annotation in OmicsBox. The bar plot shows the number of GO terms and EC numbers before and after merging eggnog based functional annotation.

A total of 44,852 genes protein coding genes were predicted in *P. alba L. “Villafranca “genome. Of these* 32,255 (71.9%) were functionally annotated (**Table** ). Support from extrinsic RNA-Seq evidence was present for the majority of predictions with 27,671 transcripts (61.7%) showing Hint-support scores > 0.4, indicating moderate to high confidence. These values are substantially higher than those previously reported for the previous *P. alba* assembly, where 32,963 genes were identified (15) and also exceeds the number protein coding genes reported for *P. tremula* (35,984) and *P. tremuloides* (36,830) (26).

**Table 3.**
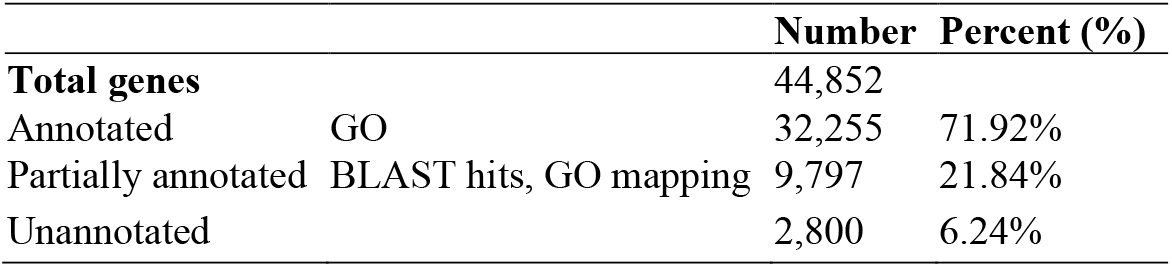
Functional annotation of predicted genes for *P. alba* L. ‘‘Villafranca” clone.

### Structural variant analysis

Genome alignment was performed with minimap2/v2.24 (29) using default parameters. Input consists of *P. alba* assembly (Query) and reference, *P. trichocarpa* genome (GCF_000002775.5). The resulting SAM file was sorted and indexed using samtools/v1.16.1 (38).

Then the SVIM-asm tool (36) was used to identify SVs (insertions, inversions, deletions, duplications, interspersed duplications and tandem duplications) from the sorted BAM file. The identified SVs were exported in VCF format for downstream analysis.

In total 9,741 SVs were detected with 3,688 insertions, with 6,038 deletions in *P. alba* relative to *P. trichocarpa*. To visualize the results Circa OM genomics software was used to (https://omgenomics.com/circa) (Figure 6).

**Figure 6.**
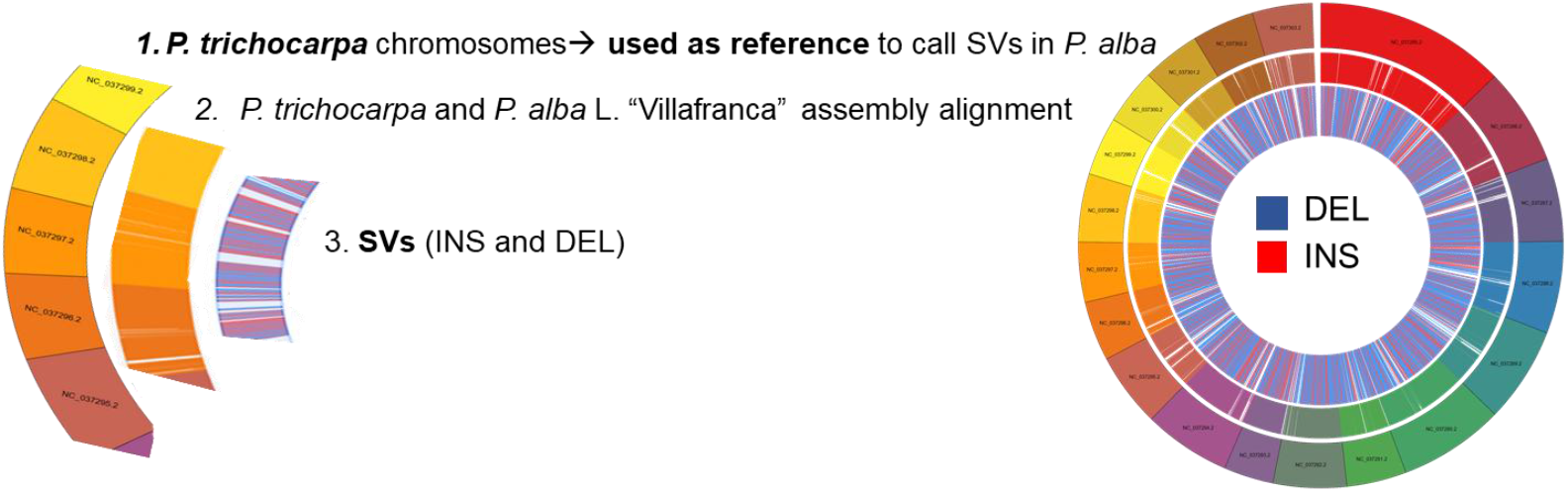
Circos plot of SVs in *P. alba* L. ‘‘Villafranca” relative to *P. trichocarpa* genome.

### Data Records

All the raw sequencing data and the genome assemblies have been submitted to NCBI. PacBio HiFi reads of *Populus alba* L. “Villafranca” NCBI are available in Sequence Read Archive under accession numbers:SRR32311034 and SRR32311033 (2025).

*Populus alba* L. “Villafranca” genome assembly data on NCBI GenBank: associated BioProject: PRJNA1214697, associated BioSample:SAMN46381866 and genome assembly accession: JBLEQM000000000

### Technical Validation

The quality and concentration of extracted DNA were assessed using Nanodrop Spectrophotometer and FEMTO size profile before the genome sequencing.

After the genome assembly was completed, the assembly results were evaluated:

i. The HiFi reads used for genome assemblies were mapped back onto to the assembled genome. The alignment rate of reads was in both cases higher than 99%, showing high consistency between the reads and assembled genomes.
ii. the completeness of the genome assemblies was evaluated using the BUSCOs embryophyta_odb10 database.

## Code availability

The manuscript did not use custom code to generate or process the data described. All data processing was performed using publicly available tools, except Figure 3 and Figure 4 were generated using GS-viewer and the complete code is available at https://github.com/mirkocelii/GS-viewer.

## Acknowledgements

This study was carried out within the Agritech National Research Center and received funding from the European Union Next-Generation EU (PIANO NAZIONALE DI RIPRESA E RESILIENZA (PNRR)– MISSIONE 4 COMPONENTE 2, INVESTIMENTO 14–DD 1032 17/06/2022, CN00000022). This manuscript reflects only the authors’ views and opinions, neither the European Union nor the European Commission can be considered responsible for them. Iqra Sarfraz Agrobioscience PhD Scholarship was funded by Programma Operativo Nazionale Ricerca e Innovazione 2014-2020 (CCI 2014IT16M2OP005), FSE REACT-EU, Azione IV.4 “Dottorati e contratti di ricerca su tematiche dell’innovazione” e Azione IV.5 “Dottorati su tematiche Green”.

## Authors contributions

I.S: Conceptualization, Methodology, Investigation, Validation, Formal analysis, Data curation, Software, Visualization, Writing-original draft, Writing-review & editing.

L.S: Conceptualization, Methodology, Resources, Formal analysis, Data curation, Software, Project administration, Funding acquisition, Writing-review & editing.

A.Z: Conceptualization, Methodology, Validation, Resources, Formal analysis, Data curation, Software, Visualization, Funding acquisition, Writing-review & editing.

A.F: Conceptualization, Writing-review & editing.

M.C: Methodology, Validation, Data curation, Software, Visualization, Writing-review & editing. R.A.W: Methodology, Resources, Writing-review & editing.

## Ethics declarations and competing interests

The authors declare no competing interests.

